# Functional genomic analysis of corals from natural CO_2_-seeps reveals core molecular responses involved in acclimatization to ocean acidification

**DOI:** 10.1101/112045

**Authors:** CD Kenkel, A Moya, J Strahl, C Humphrey, LK Bay

## Abstract

Little is known about the potential for acclimatization or adaptation of corals to ocean acidification and even less about the molecular mechanisms underpinning these processes. Here we examine global gene expression patterns in corals and their intracellular algal symbionts from two replicate population pairs in Papua New Guinea that have undergone long-term acclimatization to natural variation in pCO_2_. In the coral host, only 61 genes were differentially expressed in response to pCO_2_ environment, but the pattern of change was highly consistent between replicate populations, likely reflecting the core expression homeostasis response to ocean acidification. Functional annotations highlight lipid metabolism and a change in the stress response capacity of corals as a key part of this process. Specifically, constitutive downregulation of molecular chaperones was observed, which may impact response to combined climate-change related stressors. Elevated CO_2_ has been hypothesized to benefit photosynthetic organisms but expression changes of *in hospite Symbiodinium* in response to acidification were greater and less consistent among reef populations. This population-specific response suggests hosts may need to adapt not only to an acidified environment, but also to changes in their *Symbiodinium* populations that may not be consistent among environments. This process adds another challenging dimension to the physiological process of coping with climate change.

## INTRODUCTION

Increasing atmospheric carbon dioxide concentration contributes to global warming and alters ocean carbonate chemistry in the process known as ocean acidification (Sabine *et al*., 2004). Elevated atmospheric CO_2_ increases the hydrogen ion concentration [H+], thereby reducing ocean pH. This excess H+ reacts with carbonate ions [CO_2_^3−^] to form bicarbonate [HCO3^-^], lowering the saturation state of carbonate minerals, such as calcite and aragonite (Feely *et al.,* 2009). Many marine taxa rely on carbonate minerals to build their calcium carbonate [CaCO_3_] skeletons. Increasing H+ and concomitant reductions in pH increase the potential for dissolution of present skeletons (van Woesik *et al.,* 2013). Simultaneous reductions in the bioavailability of carbonate ions also increase the difficulty of depositing new skeleton (Kleypas *et al.,* 1999). Ocean acidification has been predicted to have major consequences for marine calcifying organisms, such as reef-building corals through this combination of effects (Hoegh-Guldberg *et al.,* 2007).

Scleractinian corals form the basis of the most biodiverse marine ecosystems on the planet: tropical coral reefs (Caley & St John, 1996, Idjada & Edmunds, 2006). They also provide important ecosystem services, such as habitat for fisheries species and shore protection (Sheppard *et al*., 2005). Consequently, investigation of coral responses to acidification has received substantial attention in recent years. The majority of empirical work has focused on relatively short-term (days to months) exposure of corals to simulated acidification in aquaria and the reported fitness consequences have been mixed. A recent meta-analysis found that for every unit decrease in the saturation state of aragonite, coral calcification declines by 15% on average, though individual studies report more significant declines or even increases (Chan & Connolly, 2013), which may be attributable to differences in tolerance among species (Albright, 2011, Erez *et al*., 2011,Jokiel, 2011).

Natural CO2-seep environments provide an attractive alternative to aquarium-based experiments aimed at understanding coral resilience potential: no experimental manipulations are necessary and *in situ* populations have likely already undergone some level of acclimatization or adaptation to be able to inhabit low-pH environments. Work by Fabricius *et al.* (2011) on corals at volcanic CO_2_-seeps in Papua New Guinea (PNG) has provided support for the mixed effects observed in laboratory experiments. Naturally acidified environments drastically alter the coral community, but some species, like massive *Porites,* appear unaffected, while others, such as Acroporids, are significantly less common or even absent (Fabricius *et al*., 2014). Population reductions *in situ,* combined with observations of negative physiological impacts, including declines in calcification under elevated pCO_2_ (Strahl *et al*., 2015) strongly suggests that acidification imposes selection pressure on less resilient taxa, such as Acroporids. Consequently, *Acropora* spp. are predicted to be ecological ‘losers’ under future acidification scenarios (Schoepf *et al*., 2013). However, the fact that some *Acropora* spp. can still be found in seep environments indicates that standing genetic variation for acidification tolerance may already exist within these less resilient species, similar to recent work in analogous natural systems investigating variation in coral thermal tolerance (D'Croz & Maté 2004, Kenkel *et al.,* 2013a, Oliver & Palumbi, 2011) and its mechanistic basis (Barshis *et al.,* 2013, Dixon *et al.,* 2015, Kenkel & Matz, 2016).

Transcriptome sequencing has become a powerful tool for investigating physiological plasticity and adaptive evolution in a changing environment and can provide insight into the mechanistic basis of population-level variation (DeBiasse & Kelly, 2016). We used RNA-seq to investigate the core genomic response underpinning long-term acclimatization to acidification in *Acropora millepora* populations in the PNG seep system. In addition to significant population declines and reduced rates of net calcification at CO2-seep sites compared to paired non-impacted reefs (Fabricius *et al.,* 2014,Strahl *et al.,* 2015), coral-associated microbial communities also differ significantly in this species. In particular, *A. millepora* at seep sites exhibit a 50% reduction in symbiotic *Endozoicomonas,* a putative mutualist and generally dominant component of the coral microbiome (Morrow *et al.,* 2015,Neave *et al.,* 2017). We evaluated global gene expression profiles in adult corals and their algal endosymbionts, *Symbiodinium* spp., from replicate pairs of control and seep environments at two different reefs in the PNG system: Dobu (control pH = 8.01, 368 μatm pCO_2_; seep pH = 7.72, 998 μatm pCO_2_) and Upa-Upasina (control pH = 7.98, 346 μatm pCO_2_; seep pH = 7.81, 624 μatm pCO_2_) (Fabricius *et al.,* 2014). We interpret consistent shifts in expression among seep-site populations in the two replicate reef systems to reflect the core molecular response involved in long-term acclimatization and/or adaptation to ocean acidification.

## METHODS

### Sampling Collection and Processing

Small tips of coral branches were collected individually from 15 *A. millepora* colonies each at the CO_2_ seep and control sites of both Dobu and Upa-Upasina Reefs, Milne Bay Province, Papua New Guinea, at 3 m depth, under a research permit by the Department of Environment and Conservation of Papua New Guinea as described previously (Fabricius *et al.,* 2014,Fabricius *et al.,* 2011). Samples were snap-frozen in liquid nitrogen within minutes of collection and maintained at temperatures <−50°C until further processing.

Samples were crushed in liquid nitrogen and total RNA was extracted individually from 59 samples using a slightly modified RNAqueous kit protocol (Ambion, Life Technologies), and DNAse treated as in Kenkel *et al.* (2011). Briefly, samples homogenized in lysis buffer were centrifuged for 2 minutes at 16100 rcf to precipitate skeleton fragments and other insoluble debris and 700 μl of supernatant was used for extraction following the manufacturers' instructions, with one additional modification: in the final elution step, the same 25 μl of elution buffer was passed twice through the spin column to maximize the concentration of eluted RNA. RNA quality was assessed through gel electrophoresis and evaluated based on the presence of the ribosomal RNA bands. One μg of RNA per sample was prepared for tag-based RNA-seq as in (Lohman *et al.,* 2016,Meyer *et al.,* 2011), with modifications for sequencing on the Illumina HiSeq platform (e.g. different adapter sequences to be compatible with the different sequencing chemistry; full protocols available at: https://github.com/z0on/tag-based_RNAseq).

Noonan *et al.* (2013) demonstrated with gel-based DGGE and direct Sanger sequencing that *Symbiodinium* types do not differ between corals found in CO_2_ seep and control environments and that *Acropora millepora* host variants of clade C, closely related to C1 and C3, in the PNG seep system. To confirm this result, we mapped reads for each sample against a reference that included *A. millepora* concatenated to *Symbiodinium* clades A, B, C and D. More than 90% of *Symbiodinium* reads were assigned to clade C across all samples (Table S1). A parallel RFLP digest (Palstra, 2000, van Oppen *et al.,* 2001) of LSU types confirmed that all corals used hosted C1 (Fig. S1), however one sample from the Dobu CO_2_- seep also appeared to have some amplifiable level of D-type symbionts, therefore to be conservative, this sample was discarded from the *Symbiodinium* expression analysis dataset.

### Bioinformatic Processing

A total of 59 libraries were sequenced on two lanes of the Illumina HiSeq2500 at the University of Texas at Austin Genome Sequencing and Analysis Facility. On average, 5.4 million sequences were generated per library (range: 2.5-16.3 million), for a total of 316.8 million raw reads. A custom perl script was used to discard duplicate reads sharing the same degenerate primer (i.e. PCR duplicates) and trim the 5’-Illumina leader sequence from remaining reads. The *fastx_toolkit* (http://hannonlab.cshl.edu/fastx_toolkit) was used to remove additional reads with a homo-polymer run of ‘A’ ≥ 8 bases, retain reads with minimum sequence length of 20 bases, and quality filter, requiring PHRED quality of at least 20 over 90% of the sequence. *Bowtie 2* (Langmead & Salzbert, 2012) was used to map filtered reads to a combined transcriptome reference: a concatenated *Acropora millepora* reference transcriptome (Moya *et al*., 2012b) and a *Symbiodinium* Clade C reference transcriptome (Ladner *et al*., 2012). Read counts were assembled by isogroup (i.e. groups of sequences putatively originating from the same gene, or with sufficiently high sequence similarity to justify the assumption that they serve the same function) for both the host and symbiont transcriptomes using a custom perl script, discarding reads mapping equally well to multiple isogroups (Dixon *et al.,* 2015). For the host transcriptome, on average, 811,704 reads per library (range: 414,605 – 2,102,534) were mapped to 45,442 unique isogroups. For the symbiont transcriptome, 277,517 reads per library (range: 96,025 – 571,019) were mapped to 24,076 unique isogroups.

### Statistical Analyses

Analyses were carried out in the R statistical environment (R Development Core Team 2013). Outlier analyses were conducted using the package *arrayQualityMetrics* (Kauffmann *et al.,* 2009). Four outliers were identified in the coral host dataset, while only one was detected in the symbiont dataset. All outlier samples were discarded. Count data for the remaining host samples (Dobu-Seep = 14, Dobu-Control = 14, Upa-Upasina-Seep = 14, Upa-Upasina-Control = 13) and symbiont samples (Dobu-Seep = 14, Dobu-Control = 15, Upa-Upasina-Seep = 14, Upa-Upasina-Control = 15) were analyzed using the package *DESeq* (Anders & Huber, 2010). Dispersion estimates of raw counts were obtained by maximizing a Cox-Reid adjusted profile likelihood of a model specifying population origin and seep environment for each sample and the empirical dispersion value was retained for each gene. Low-expression genes were excluded from subsequent analyses by removing isogroups with read count standard deviations in the bottom 60% quantile of both datasets, which were identified as the filter statistics best satisfying the assumptions of independent filtering as implemented in the package *genefilter* (Gentleman *et al.*). This left 18,177 highly expressed isogroups in the coral host dataset and 9,629 isogroups in the symbiont dataset. In each dataset, expression differences were evaluated with respect to reef site (Upa-Upasina/Dobu), and pCO_2_ environment (Seep/Control) and the interaction using a series of generalized linear models implemented in the function *fitNbinomGLMs*. Multiple test correction was applied using the method of Benjamini and Hochberg (1995). Analyses were also repeated independently for each population to verify candidate gene significance with respect to seep environment.

Functional enrichment analyses were conducted using the package GO-MWU (Voolstra *et al*., 2011) to identify over-represented gene ontology (GO) terms with respect to origin and seep environment using both the classical categorical test and a rank-based methodology (Dixon 2015). The package *made4* (Culhane *et al*., 2005) was used to conduct a between-groups analysis of seep and control samples within each dataset to identify the most discriminatory genes in terms of differential expression between reef environments. A permutation test was used to evaluate whether there were significantly more differentially expressed genes in the symbiont dataset relative to the coral host dataset. Since FDR-correction is partially based on the number of tests conducted, we created 1,000 random 9,629 gene subsets of the host 18,177 gene dataset and repeated FDR-correction on this reduced sample. We then compared the distribution of significant tests obtained in the subsample to the observed symbiont gene set to obtain an estimate of significance.

## RESULTS

In total, 571 isogroups (genes) were differentially expressed at the FDR cut-off level P_adj_ < 0.1 in the coral host (3% of total, Fig. 1a). The grand majority of these differences were due to reef origin (Dobu *vs*. Upa-Upasina, 503 genes, Table S2). Only 61 genes were differentially regulated between corals originating from control and seep environments, 53 of which exhibited consistent differences irrespective of reef origin (Fig. 1a,c, Table S3). Significantly more expression changes were detected in *Symbiodinium* populations (P_permutation_<0.0001) where a total of 1123 genes were differentially expressed (P_adj_ < 0.1, 12% of total, Fig 1b). Again, the majority of these changes were attributable to differences in reef origin (Table S4), but 201 genes exhibited altered expression in seep environments relative to controls (Fig. 1b, Table S5). Expression changes in symbionts were also less consistent between populations (Fig. 1d). The purpose of this study was to evaluate expression differences following lifelong acclimatization to elevated pCO_2_ in corals. Therefore we focus on genes regulated with respect to seep environment, although differential expression patterns for genes responding to reef origin and associated functional enrichments can be found in the supplementary material (Tables S2, S4, Fig S2).

**Figure 1.**
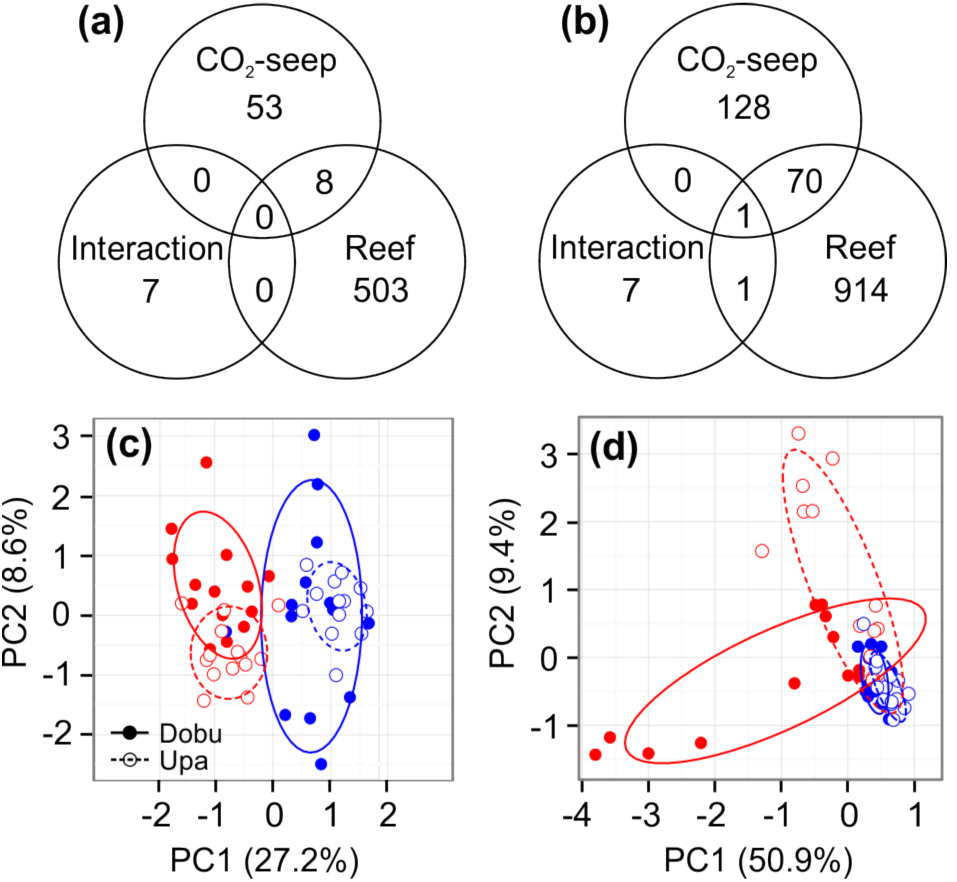
Venn diagrams of differentially expressed genes by factor (FDR-adjusted P<0.1) for host (a) and symbiont (b). Principal components analysis of top 50 most significantly differentially expressed genes by CO2-seep (red=seep, blue=control) and reef origin for host (c) and symbiont (d).

## Differential expression of coral host genes by CO_2_ seep environment

Of the 61 genes showing common population-level responses to the CO_2_-seep environment, 53 exhibited consistent baseline expression levels between corals from the different reef locations (Fig. 1a, ‘CO_2_ seep’). Of these, 26 were upregulated and 27 were downregulated in CO_2_-seep environments. Roughly half (51%) of these genes have no annotation, and thus their functions cannot be determined. We report expression patterns among annotated candidates only but the data for all differentially expressed genes can be found in Table S3. We first consider individual candidate genes and then describe altered functional processes identified through enrichment analyses.

## De novo *candidate genes*

Among annotated genes significantly upregulated in seep-site corals, three associated with transcriptional regulation were also identified in a between-groups analysis as the most discriminatory genes between seep and control samples (Fig. 2a). Two are transcriptional regulators (ig19425, ig12770, 1.08-fold and 1.09-fold, respectively) and the third is a transcription factor in the basic leucine-zipper superfamily (ig10473, 1.2-fold). In the entire *A. millepora* transcriptome, 26 genes are annotated as ‘transcriptional regulators’ and another 8 are bZIP transcription factors. A methyl-CpG binding transcriptional regulator (ig9532) was also upregulated by 1.05-fold in corals from seep sites, but showed an additional effect of host origin, with corals from Dobu having higher baseline expression than corals from Upa-Upasina (Fig. 2c). This methyl-CpG binding regulator was one of only three genes with this annotation in the entire *A. millepora* transcriptome, the other two of which (ig16785 and ig21898) were not found in the final expression set. A transcriptional repressor in the hairy/E (spl) family (ig7904) was among the most discriminatory genes and down-regulated in response to seep environments by 1.16-fold (Fig. 2b), again suggesting some role for transcriptional regulation, though 13 isogroups in the transcriptome also have this same annotation.

**Figure 2.**
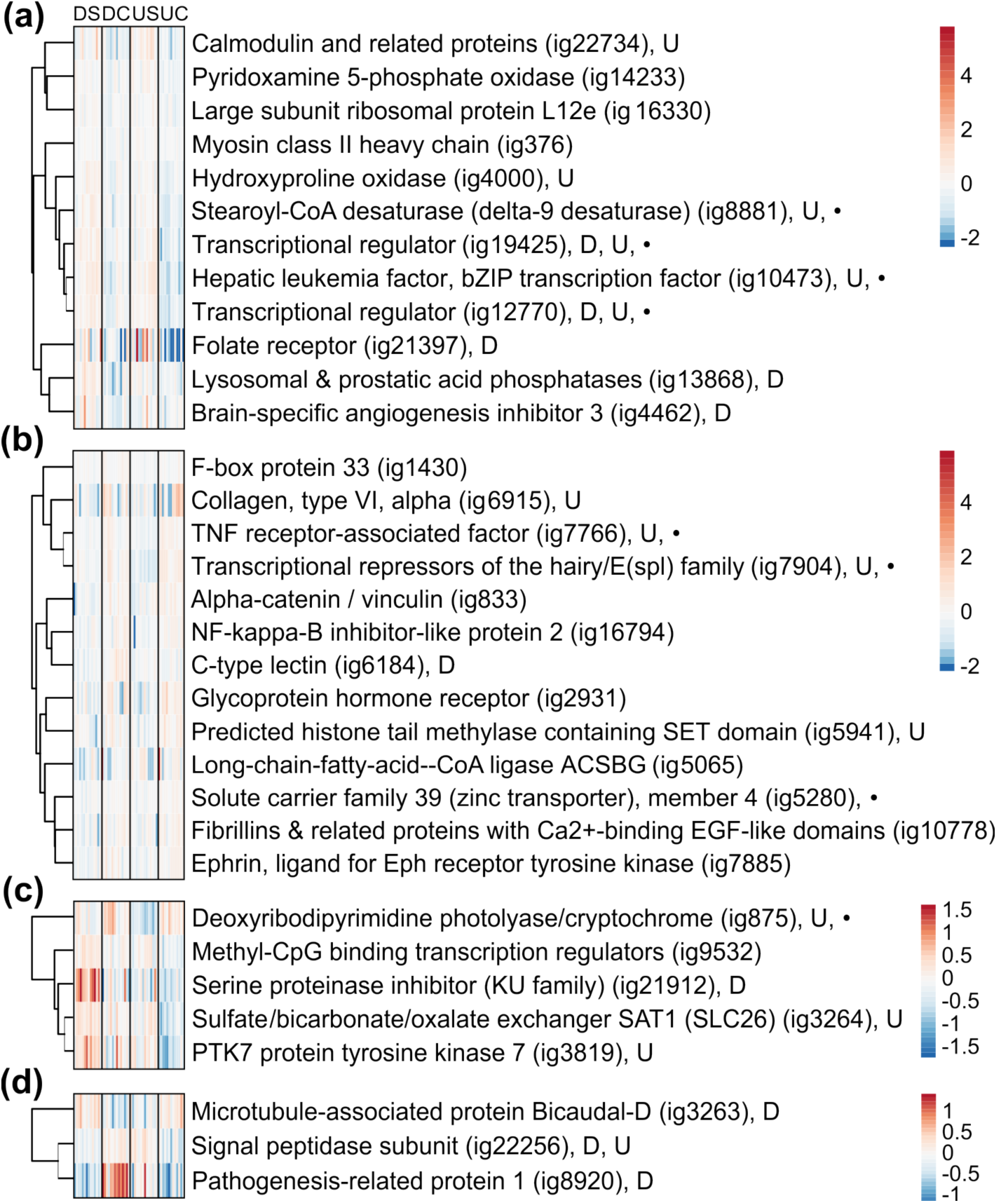
Heatmaps of annotated genes (FDR-adjusted P<0.1) in the coral host that showed upregulation in response to seep environment (a), downregulation in response to seep environment (b), an effect of reef origin in addition to an effect of seep environment (c) or a reef origin x seep environment interaction (d). D=FDR-adjusted P<0.1 in Dobu-only dataset; U=FDR-adjusted P<0.1 in Upa-Upasina-only dataset; •=Top discriminatory gene as identified via between-groups analysis for seep environment. DS=Dobu-seep, DC=Dobu-control, US=Upa-Upasina-seep, UC=Upa-Upasina-control.

A TNF receptor-associated factor (ig7766) is also a top discriminatory gene. This family, involved in the innate immune response, recently came to prominence given its putative role in the coral stress response (Barshis *et al*., 2013). Its downregulation, together with an NF-kappa-B inhibitor (ig16794) and a c-type lectin (ig6184, Fig. 2b), highlight a potential impact of elevated pCO_2_ on the innate immune response. However, 104, 49 and 143 isogroups respectively have identical annotations in the *A. millepora* transcriptome.

An alpha-catenin/vinculin isoform (ig833), one of three genes with this annotation, is downregulated in seep site corals by 1.15-fold (Fig. 2b). The other two isoforms (ig1210 and ig21857) are not differentially expressed and not included in this expression dataset. Additional cytoskeletal components including a collagen (ig6915) and fibrillin (ig10778) are also downregulated by 1.23 and 1.17-fold respectively, although these annotations are fairly common (86 and 372 isogroups in the transcriptome, respectively).

## Categorical Functional Enrichments

A categorical functional enrichment analysis did not reveal any statistically significant candidates following FDR-correction. The top three ‘biological process’ enrichments were ‘small molecule biosynthetic process’ (GO:0044283, P_Raw_ = 0.1), ‘fatty acid metabolic process’ (GO:0006631, P_Raw_ = 0.3) and ‘small molecule catabolic process’ (GO:0044282, P_Raw_ = 0.3), which resulted from a set of four candidate genes. Pyridoxamine 5-phosphate oxidase (ig14233, upregulated by 1.06-fold in seep-site corals, GO:0044283, Fig. 2a), an enzyme catalyzing the rate-limiting step in vitamin B_6_ metabolism is an annotation only assigned to one other gene in the host transcriptome (ig27779) that was not differentially expressed with respect to either seep environment or reef origin.

Hydroxyproline oxidase (ig4000, GO:0044283, GO:0044282, Fig 2a), hypothesized to play a role in activation of the apoptotic cascade (Cooper *et al*., 2008), is also upregulated by 1.07- fold in seep site corals. The only other gene of the transcriptome with this annotation (ig1278) is differentially regulated with respect to reef origin, showing 1-fold upregulation in corals from Dobu (P_Reef_ < 0.1, Table S2).

The remaining two genes are primarily involved in fatty-acid metabolism. Stearoyl-CoA desaturase (ig8881, GO:0044283, GO:0006631) is upregulated in seep sites by 1.15– fold. There are only 5 isogroups in the transcriptome with this annotation, 3 occur in the final expression list, but this isoform is the only one differentially expressed. The other candidate, long-chain-fatty-acid—CoA ligase, or long-chain acyl–CoA synthetase (ig5065, GO:0006631, GO:0044282), is downregulated by 1.19-fold and is one of only seven isoforms with this annotation. One other isoform is differentially expressed with respect to reef origin, with greater expression in Dobu-origin corals (ig3997, P_Reef_ < 0.1, Table S2), but remaining isoforms (ig2622, ig2781, ig5009, ig5135, ig12633) were not differentially expressed.

## Rank-based Functional Enrichments

Given the low number of candidate genes that passed the FDR threshold, a rank-based methodology was also used to determine functional enrichments among generally upregulated (red) and downregulated (blue) ontologies in corals from CO_2_-seep environments (Fig. 3a). ‘Chaperone-mediated protein folding’ (GO:0061077) was the top enrichment among genes downregulated in CO_2_-seep sites (Fig 3a, b). ‘Ribonucleoprotein complex biogenesis’ and ‘one-carbon metabolic process’ were the top two most enriched functional ontologies among genes upregulated in seep sites (GO:0022613 and GO: 0006730, respectively, Fig. 3 a, c, d). Interestingly, the most significantly differentially regulated genes within ‘one-carbon metabolic process’ are all individually annotated as carbonic anhydrases (Fig 3c).

**Figure 3.**
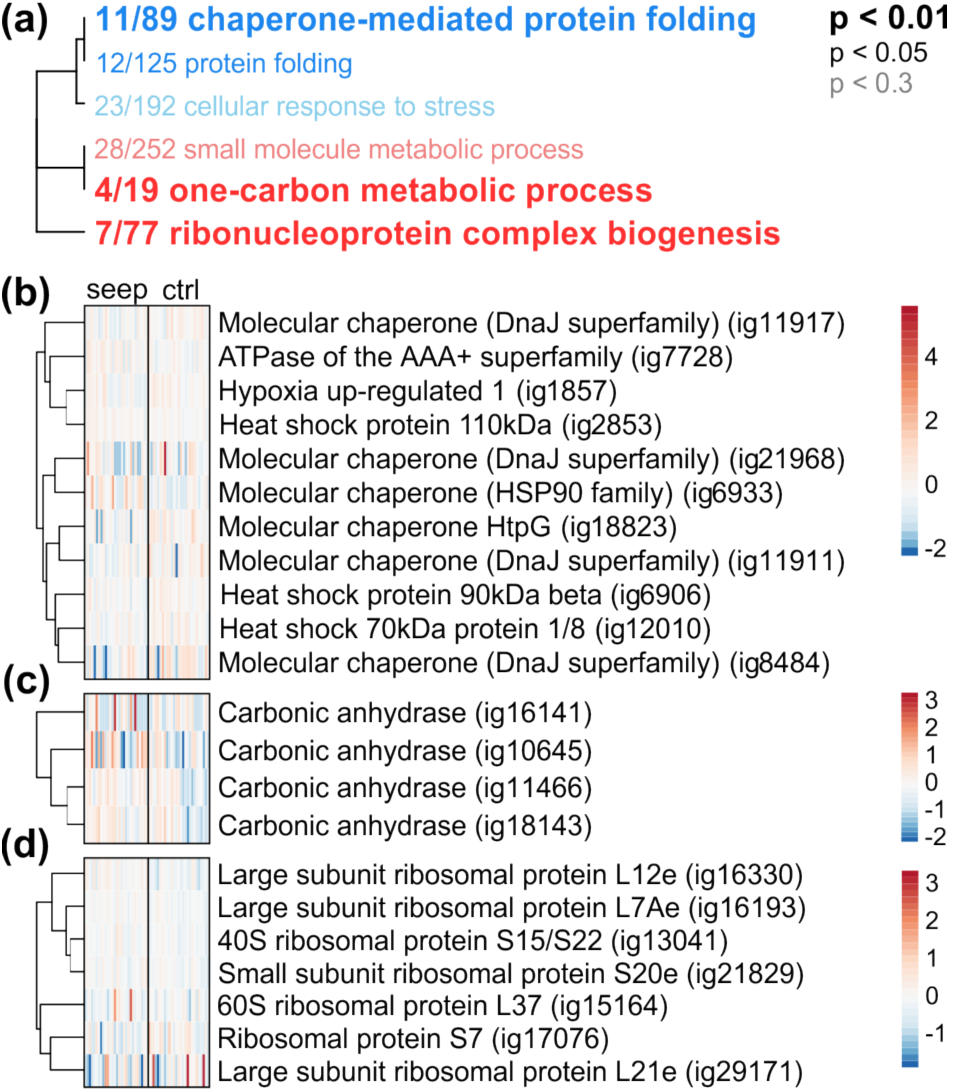
Hierarchical clustering of enriched gene ontology terms (‘biological process’) among upregulated (red) and downregulated (blue) genes in the coral host with respect to CO_2_-seep (a). Font indicates level of statistical significance (FDR-corrected). Term names are preceded by fraction indicating number of individual genes within each term differentially regulated with respect to seep site (unadjusted P<0.05). Heatmaps of these ‘good gene’ fractions are shown for ‘chaperone-mediated protein folding’ (b), ‘one-carbon metabolic process’ (c) and ‘ribonucleoprotein complex biogenesis’ (d).

## Differential expression of Symbiodinium genes by CO _2_ seep environment

Of the 201 genes differentially expressed in response to CO_2_-seep environment, 128 exhibited similar baseline expression levels between symbionts in corals from the different reef locations (Fig. 1b, ‘CO_2_ seep’, Table S5). Of these, 96 were upregulated and 32 were downregulated in CO_2_-seep environments. Only 40% of these genes were annotated, and we again report expression patterns among these candidates only, although the data for all differentially expressed genes can be found in Table S5. To enhance the sparse knowledge on *Symbiodinium* responses to acidification, we report altered functional processes identified through categorical and rank-based enrichment analyses.

### Categorical Functional Enrichments

The relatively small number of genes responding to seep site and a lack of annotations resulted in no statistically significant ontology terms following FDR-correction of a categorical enrichment analysis. The top three ‘biological process’ enrichments were ‘regulation of chromosome organization’ (GO:0033044, P_Raw_ = 0.005), ‘response to bacterium’ (GO:0009617, P_Raw_ = 0.03) and ‘regulation of organelle organization’ (GO:0033043, P_Raw_=0.05), which resulted from a set of six candidate genes. A peptidyl-prolyl cis-trans isomerase in the Ess family, matching Ess1 (c78122, GO:0033044, GO:0033043 Fig. 4a) is upregulated by 1.05-fold at seep sites. In the *Symbiodinium* Clade C transcriptome 62 clusters are annotated as PPIs, which catalyze the *cis–trans* isomerisation of peptide bonds N-terminal to proline residues in polypeptide chains, but this is the only cluster to have homology with Ess1. An E3 SUMO-protein ligase pli1 (c28523, GO:0033044, GO:0033043, Fig 4c) was also upregulated in seep site corals by 1.1-fold, but shows an additional effect of reef origin, with *Symbiodinium* in Dobu corals having higher baseline expression than *Symbiodinium* in Upa-Upasina corals. This annotation occurred twice in the transcriptome, but the other gene (c71663) was not included in the final expression set. The third gene in this regulatory group, the meiosis protein mei2 (c26263, GO:0033043, Fig 4a) was also upregulated by 1.1-fold at seep sites. Three other clusters in the transcriptome were assigned this annotation (c_sym_78605, c49233_81271, c94595), two of which were in the final expression set analyzed here and one was significantly differentially expressed with respect to reef origin (c94595, 1.02-fold, Table S3).

**Figure 4.**
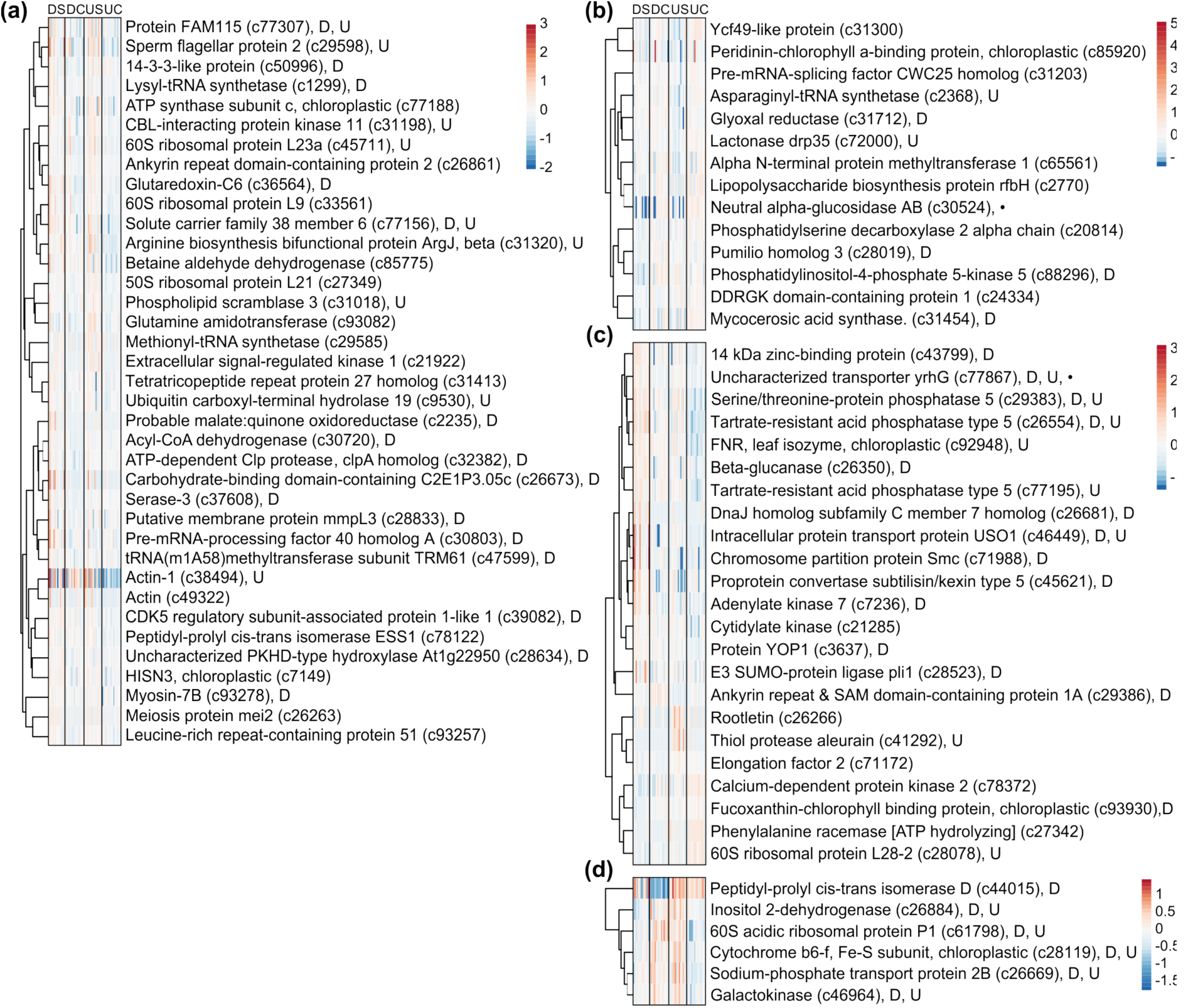
Heatmap of annotated genes (FDR-adjusted P<0.1) in *Symbiodinium* that showed upregulation in response to seep environment (a), downregulation in response to seep environment (b), an effect of reef origin in addition to an effect of seep environment (c) or a reef origin x seep environment interaction (d). D=adjusted P<0.1 in Dobu-only dataset; U=adjusted P<0.1 in Upa-Upasina-only dataset; •=Top discriminatory gene as identified via between-groups analysis for seep environment. DS=Dobu-seep, DC=Dobu-control, US=Upa-Upasina-seep, UC=Upa-Upasina-control.

The last three genes were all involved in bacterial response (GO:0009617) and all upregulated in seep sites. One of the genes was a 14-3-3-like protein (c50996, 1.06-fold, 9 genes with this annotation in transcriptome, Fig. 4a). The second was an ankyrin repeat domain-containing protein 2 (c26861, 1.02-fold, 9 genes with this annotation, Fig. 4a). The last was tartrate resistant acid phosphatase type 5 (c26554, 1.1-fold, Fig 4c, 11 genes with this annotation) and also showed an effect of origin, with *Symbiodinium* in Dobu corals having higher baseline expression than *Symbiodinium* in Upa-Upasina corals.

### Rank-based Functional Enrichments

The only significant functional enrichment identified with rank-based analysis was ‘translation’ (GO:0006412, Fig. 5a) which was enriched among genes upregulated in seep sites. Individual genes within this term were primarily annotated as ribosomal proteins (Fig. 5b).

**Figure 5.**
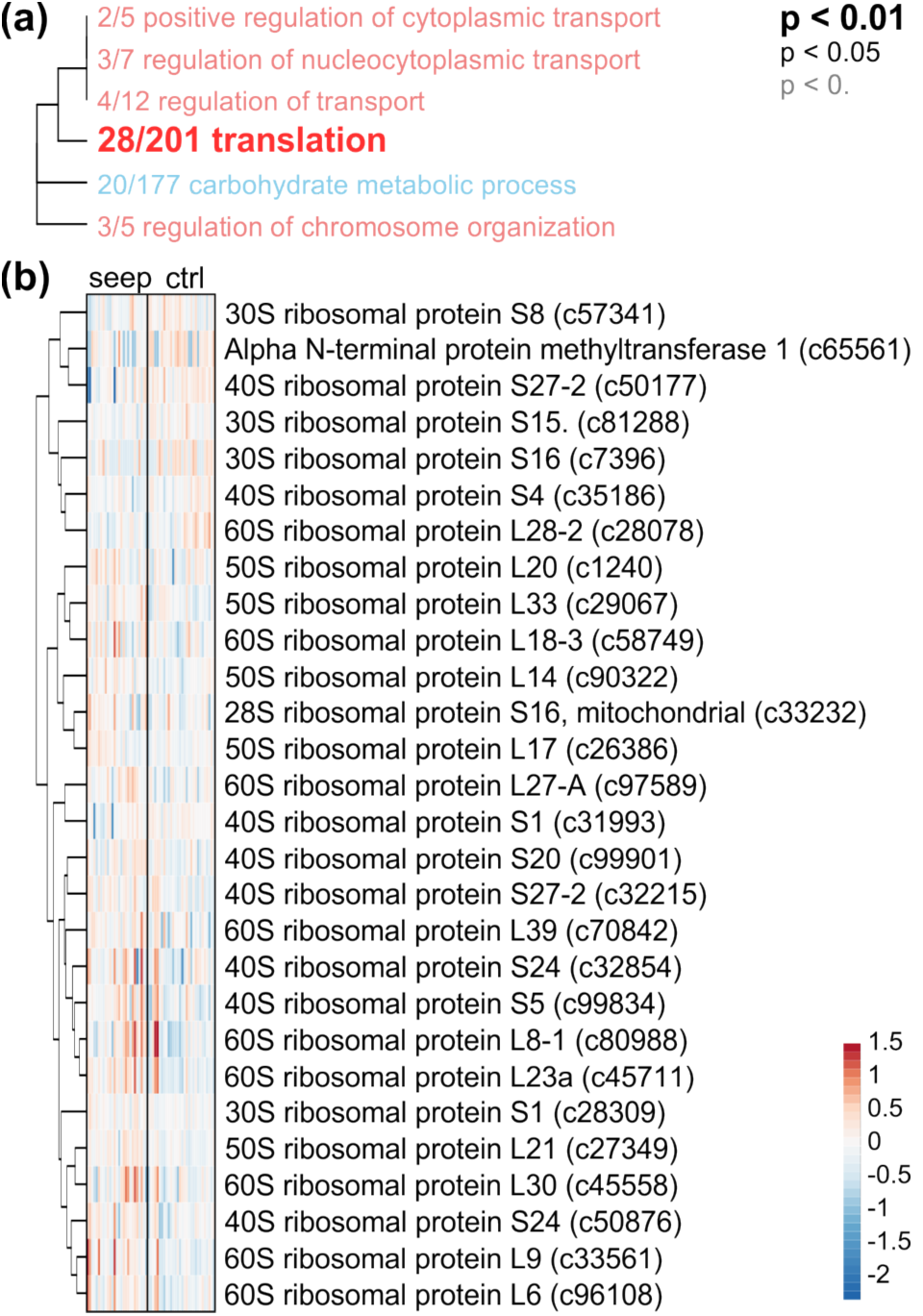
Hierarchical clustering of enriched gene ontology terms (‘biological process’) among upregulated (red) and downregulated (blue) symbiont genes with respect to CO_2_-seep (a). Font indicates level of statistical significance (FDR-corrected). Term names are preceded by fraction indicating number of individual genes within each term differentially regulated with respect to seep site (unadjusted P<0.05). A heatmap of this ‘good gene’ fraction is shown for ‘translation’ (b).

## DISCUSSION

The aim of this study was to investigate the genomic basis of acclimatization to chronic exposure to ocean acidification in a reef-building coral through a comparison of closely situated control and CO_2_ impacted sites (500 m and 2500 m at Upa-Upasina and Dobu, respectively, with 30 km between the two populations). Uniquely, we report global gene expression profiles in both the coral host and their *in hospite Symbiodinium* that have undergone life-long acclimatization to naturally acidified environments. Previous population level studies (Fabricius *et al*., 2014, Morrow *et al*., 2015,Strahl *et al*., 2015) strongly suggest that acidified environments impact the fitness of *Acropora millepora.* Despite this, we found very few consistent changes in global gene expression patterns between control and seep sites (Fig. 1). This may be because gene expression changes did not reflect actual protein content or because of post-translational regulation (Greenbaum *et al*., 2003). It is also possible that substantial inter-individual variation in expression (e.g.Bay *et al*., 2009,Csaszar *et al*., 2009) masked the detectability of expression differences in response to environmental pCO_2_. On the other hand, important biochemical health measures related to cell protection and cell damage were unaffected in *A. millepora* in response to elevated pCO_2_ up to 800 μatm at the same sites studied here (Strahl et al. 2015), consistent with our findings of a minimal expression response.

The absence of significant gene expression changes may not necessarily be surprising if acidification is a chronic stressor for the corals. Cellular stress gene expression responses are transient and non-specific (Kültz, 2005). Once immediate damage is repaired, a secondary, permanent cellular homeostasis response occurs, which is specific to the triggering stressor and which facilitates the maintenance of homeostasis under the new environmental regime (Kültz, 2003). It is likely that *A. millepora* exhibits open populations in this system given the broadcast spawning behavior of this species and the close proximity of study sites. Newly recruited juvenile corals may have exhibited an initial stress response, but their gene expression baselines shifted with age in order to acclimate to their local environment. Moya *et al.* (2015) previously reported dampened expression responses in a time-series exposure of juvenile *A. millepora* to elevated pCO_2_, consistent with this hypothesis. Therefore, the small but constitutive differences in expression detected here, in two replicate populations (n = 13 – 15 colonies per site) acclimatized to CO_2_-seep environments, likely reflects the core expression homeostasis response to ocean acidification.

In the coral host, this core response involves changes in gene regulation involved in fatty acid (FA) metabolism (Fig. 2). Differential regulation of stress response genes also occurred: specifically, corals from seep environments constitutively down-regulate expression of molecular chaperones (Fig 3a, b). Interestingly, we did not find explicit signatures indicating altered expression in calcification related genes, though some carbonic anhydrase isoforms did appear to be constitutively upregulated in seep environments (Fig 3c), but those could also be involved in cellular pH homeostasis. Finally, expression changes in *in hospite Symbiodinium* were greater, and unlike patterns in their coral hosts, were not consistent in seep habitats among reefs (Fig. 1c, d), which may have implications for the symbiosis.

### Differential regulation of fatty acid metabolism

The combined upregulation of a FA synthesis gene (Stearoyl-CoA desaturase) and downregulation of a FA catabolism gene (Long-chain-fatty-acid-CoA ligase), both key enzymes in their respective functional pathways (Dobrzyn *et al*., 2004,Watkins, 1997) and fairly unique annotations within the *A. millepora* transcriptome, suggest that coral lipid metabolism is modified in the process of acclimatization to acidification. Recent work on the transcriptomic response of urchins (*Strongylocentrotus purpuratus*) to experimental ocean acidification found that populations which naturally experience more frequent low pH conditions also differentially regulated fatty acid metabolic pathways (Evans *et al*., 2017). Interestingly, differential regulation of lipid metabolism genes was also observed in prior laboratory experiments on corals exposed to acute acidification stress (Moya *et al*., 2012b), but this particular functional process was not specifically discussed. Eleven clusters encoding fatty acid synthases were found in the *A. millepora* transcriptome, and most of them were upregulated in response to acute acidification stress (A. Moya unpublished data).

Our results indicate a metabolic shift in CO_2_-seep site corals in favor of increasing lipid storage. This is supported by findings of Strahl et al. (2015), who detected slightly higher ratios of storage to structural lipids in *A. millepora* at seep *vs.* control sites at Dobu and Upa-Upasina. Stearoyl-CoA desaturase catalyzes the rate-limiting step in the synthesis of unsaturated fatty acids, which are components of both structural (e.g. membrane phospholipids) and storage lipids (e.g. triacylglycerol, wax esters, sterol ester), and the disruption of genes encoding this enzyme in mice leads to reduced body adiposity (Ntambi *et al*., 2002). Long-chain-fatty-acid-CoA ligase, on the other hand, activates the first step of fatty acid metabolism or β-oxidation (Watkins, 1997), when lipids are being broken down. Storage lipids such as wax esters, triacylglycerol and free fatty acids are critical components of corals’ energetic status (Edmunds & Davies, 1986,Harland *et al*., 1993) and depletions in lipid stores can impact long-term survival and reproduction (Anthony *et al*., 2009). Furthermore, genes involved in lipid metabolism were found to exhibit significantly elevated rates of protein evolution in Acroporids, but the authors were unable to speculate about putative adaptive roles for lipid metabolism (Voolstra *et al*., 2011).

Recently, Strahl *et al.* (2016) found that *A. millepora* from the Dobu seep site tend to have elevated levels of total lipid and protein, as well as elevated levels of FAs (including polyunsaturated FA) relative to control site corals, in support of observed expression differences. Other studies have also found significant changes in lipid content in response to acidification. In two separate aquarium-based acidification experiments, lipid content in *A. millepora* was found to increase following exposure to elevated pCO_2_ (Kaniewska *et al*., 2015,Schoepf *et al*., 2013). Behavioral changes may also be involved in this pattern as both feeding rate and lipid storage increased in *Acropora cervicornis* under simulated acidification (Towle *et al*., 2015). Whether the mechanism is behavioral plasticity or adaptive genetic change in lipid metabolic capacity, the combined evidence suggests that lipid metabolism likely plays a role in a coral’s capacity to withstand ocean acidification and future work should aim to investigate the mechanistic basis of this process.

### Downregulation of chaperones

Upregulation of chaperones is a hallmark of the acute cellular stress response (Gasch *et al*., 2000), but is usually transient as constitutive upregulation of heat shock proteins is costly and can result in decreased growth and fecundity (Sørensen *et al*., 2003). In *Drosophila* and soil isopods exposed to chronic stress, Hsp70 expression is reduced rather than elevated (Köhler & Eckwert, 1997,Sørensen *et al*., 1999). HSPs are also known to be constitutively downregulated following long-term thermal stress in corals (Kenkel *et al*., 2013b, Meyer *et al*., 2011,Sharp *et al.,* 1997). Short-term laboratory manipulations suggest that exposure to acidification prompts expression of immediate stress response genes, like HSPs (Moya *et al*., 2012b,Moya *et al*., 2015); and Kaniewska *et al* (2012) observed downregulation of chaperones following one month of elevated pCO_2_ exposure. Therefore, acute exposure to acidification conditions is stressful for *A. millepora* and the constitutive downregulation of HSPs observed here is likely a consequence of chronic exposure to elevated pCO_2_ at the seep sites.

Given that HSP induction is critical for mounting a successful thermal stress response, the suppression of baseline HSP expression levels induced by acidified environments may impact the capacity of *A. millepora* to cope with the synergistic effects of global climate change. Acidification is predicted to become a chronic stress on reefs worldwide if climates continue to change (Hoegh-Guldberg *et al*., 2007). While temperatures will simultaneously increase, extreme thermal anomalies are also predicted to become more frequent and severe (Frich *et al*., 2002). Our results suggest that the combined effects of acidification and temperature stress may be more detrimental than acidification alone because of the dampening effects of chronic exposure on the cellular stress response. Some laboratory manipulations have found synergistic negative impacts of combined acidification and temperature; for example, calcification of *Stylophora pistillata* decreased by 50% under both elevated temperature and pCO2, but was unchanged under each stressor individually (Reynaud *et al*., 2003). However, bleaching surveys following a minor thermal stress event in PNG did not indicate that acidified reefs suffered increased bleaching relative to control reefs (Noonan & Fabricius, 2015). It will be critical to determine whether constitutive downregulation of HSPs resulting from long-term pCO_2_ exposure makes it more difficult for a coral to subsequently upregulate HSPs to counter acute thermal stress, or if other mechanisms or isoforms are employed to counter acute thermal stress in chronically acidified environments.

### No significant differential regulation of calcification genes

We did not observe functional enrichments indicating differential regulation of calcification related genes overall in *A. millepora,* although some carbonic anhydrase isoforms were constitutively upregulated in seep site corals (Fig. 3c). Experimentally, some coral species have been shown to maintain (Reynaud *et al*., 2003) and even increase (Castillo *et al*., 2014) calcification during laboratory acidification experiments and this effect has been hypothesized to result from the ability of corals to alter carbonate chemistry at the site of calcification (McCulloch *et al*., 2012,Venn *et al*., 2013). Our *a priori* expectation was that expression patterns of calcification related genes should be altered to affect this physiological rescue. *Pocillopora damicornis* were observed to upregulate HCO_3_- transporters at moderately low pH (7.8 and 7.4;Vidal-Dupiol *et al*., 2013), while *Siderastrea sidereal* upregulated expression of H^+^ ion transporters (Davies *et al*., 2016) consistent with this hypothesis.

However, *A. millepora* does not appear to conform to this expectation. Expression of calcification related genes significantly changed in *A. millepora* following short-term 3-day acidification stress exposure (Moya *et al*., 2012b), but these effects dissipate when experimental treatment periods are extended (Kaniewska *et al*., 2012, Moya *et al*., 2015,Rocker *et al*., 2015; 28, 9 and 14 days, respectively). Furthermore, *A. millepora* from PNG seep sites had reduced levels of net calcification, resulting from decreases in dark calcification, compared to neighboring control reef sites (Strahl *et al*., 2015). This suggests that *A. millepora* has a reduced capacity to actively alter pH at the site of calcification in the absence of additional photosynthetic energy (i.e. in the dark,Strahl *et al*., 2015). The regulation of cellular pH at calcification sites is an energetically costly process (Al-Horani, 2005, Barnes & Chalker, 1988). Given that calcification related gene expression is plastic in *A. millepora* on shorter time-scales (Moya *et al*., 2012a), it is possible that the lack of constitutive differential regulation under long-term acidification, and subsequent decrease in net calcification, were not necessarily due to a lack of genetic variation in the ability to actively regulate these genes, but a result of trade-offs in allocation of finite energetic resources to other less costly processes that maximize net fitness under acidification stress. Indeed, Strahl *et al.* (2016) hypothesized that *A. millepora* may invest in increased tissue biomass rather than skeletal growth under acidified conditions based on prior experimental observations of unchanged or increased biomass in combination with reduced calcification (Krief *et al*., 2010, Schoepf *et al*., 2013,Strahl *et al*., 2015).

### Inconsistent changes in Symbiodinium expression profiles

More significant differences in gene expression were detected for *Symbiodinium* than for host corals between control and elevated pCO2 sites examined here. This corroborates findings from *Pocillopora damicornis* where their *in hospite* clade C *Symbiodinium,* symbionts demonstrated a more pronounced expression response following a 2-week exposure to elevated temperature, although this difference was no longer evident after 36- weeks (Mayfield *et al*., 2014). Kaniewska *et al.* (2015) examined metatranscriptomic expression responses of coral holobionts to future climate change scenarios, but their analysis method did not explicitly compare host and symbiont. Kenkel and Matz (2016) reported expression of both host and symbionts in *P. astreoides* corals reciprocally transplanted between reef habitats, but again, their network-based analytical approach precludes a direct comparison with results uncovered here. A reanalysis of their dataset with the method used here found that 14.8% of the host transcriptome was significantly differentially expressed with respect to transplant environment, while only 1.4% of the symbiont transcriptome was altered (Kenkel, unpublished data). Given the paucity of studies examining global expression of both partners in response to environmental stress (to our knowledge, the present study is only to examine expression under elevated pCO_2_), it is difficult to draw any conclusions regarding the present patterns. More data are needed to determine whether there are any consistent patterns in *Symbiodinium* gene expression responses relative to those of their host corals.

Expression changes of *in hospite Symbiodinium* showed greater differences between control and seep sites across reefs compared to the coral host (Fig. 1c vs. d). The dominant *Symbiodinium* types did not differ among corals at control and seep sites (Table S1, Fig. S1,Noonan *et al*., 2013) but it is possible that undetected background *Symbiodinium* clades or types impacted expression levels if reads from other types failed to map to the Clade C reference transcriptome used here (Ladner *et al*., 2012). However, if differences in rare *Symbiodinium* types or their expression patterns were consistent among reefs, we would still expect to observe consistent changes in expression with respect to seep environment. Conversely, if there was an interaction between potential differences in background clades or types and seep environment, this could explain the variation observed (Fig. 1d). Control site expression profiles are remarkably similar between reef sites, and the major axis of variation differentiated seep from control site populations. However, the second principal component describes variation in expression that is largely the result of divergence between seep site expression of Dobu and Upa-Upasina origin corals (Fig. 1d).

Consistent expression changes among control and seep site *Symbiodinium* implicated an alteration of the biological process of translation: specifically, many ribosomal proteins were constitutively upregulated at the seep sites (Fig. 5). Ribosome production is intimately tied to cell growth, and known to regulate cell size and the cell cycle (Jorgensen & Tyers, 2004). Net photosynthesis was significantly elevated in *A. millepora* from CO_2_–seep sites (Strahl *et al*., 2015), potentially as a result of enhanced *Symbiodinium* cell growth or division (and hence elevated expression of ribosomal proteins) although these processes remain to be quantified. Determining the mechanistic drivers of divergence between seep site populations among Dobu and Upa-Upasina reefs is more difficult. Of the top 10 gene loadings for PC2 (Fig. 1d), 8 had no annotation, precluding speculation about function. Nevertheless, the complexity of the response in *Symbiodinium* may have implications for the symbiotic interaction, if the coral host has to respond to the dual impacts of changes in its external environment, and its symbiont community. It is recognized that mutualisms are more susceptible to climate change impacts because the inherent inter-dependency between species means that even though stress only impacts one partner, both partners ultimately share the cost (Kiers *et al*., 2010). There are many knowledge gaps remaining for both major global change stressors, however, our understanding of thermal stress impacts on the coral-algal symbiosis far outstrips understanding of acidification impacts (Barshis, 2015). Filling this gap will be critical for refining predictions of coral response to continued acidification and the combined impacts of global climate change.

## DATA ARCHIVING

Raw RNA Tag-seq data have been uploaded to NCBI’s SRA: PRJNA362652. R scripts and input files for gene expression analyses will be archived on DRYAD upon manuscript acceptance. R scripts for ontology enrichment analyses and directions for formatting input files can be found at http://www.bio.utexas.edu/research/matz_lab/matzlab/Methods.html

## ACKNOWLEDGEMENTS

We thank the communities at Dobu and Upa-Upasina for their permission to study the corals on their reef. Many thanks to Katharina Fabricius, Sam Noonan, Sven Uthicke and the crew of the M.V. Chertan for their support during field work. We thank P. Davern and M. Donaldson for their help with the logistics and shipment of the equipment, and QantasLink for continued support. Catarina Schlott crushed samples for RNA extractions. Bioinformatic analyses were carried out using the computational resources of the Texas Advanced Computing Center (TACC). This project was funded by the Australian Government's National Environmental Research Program and the Australian Institute of Marine Science.

## SUPPLEMENTARY FIGURES

**Figure S1.**
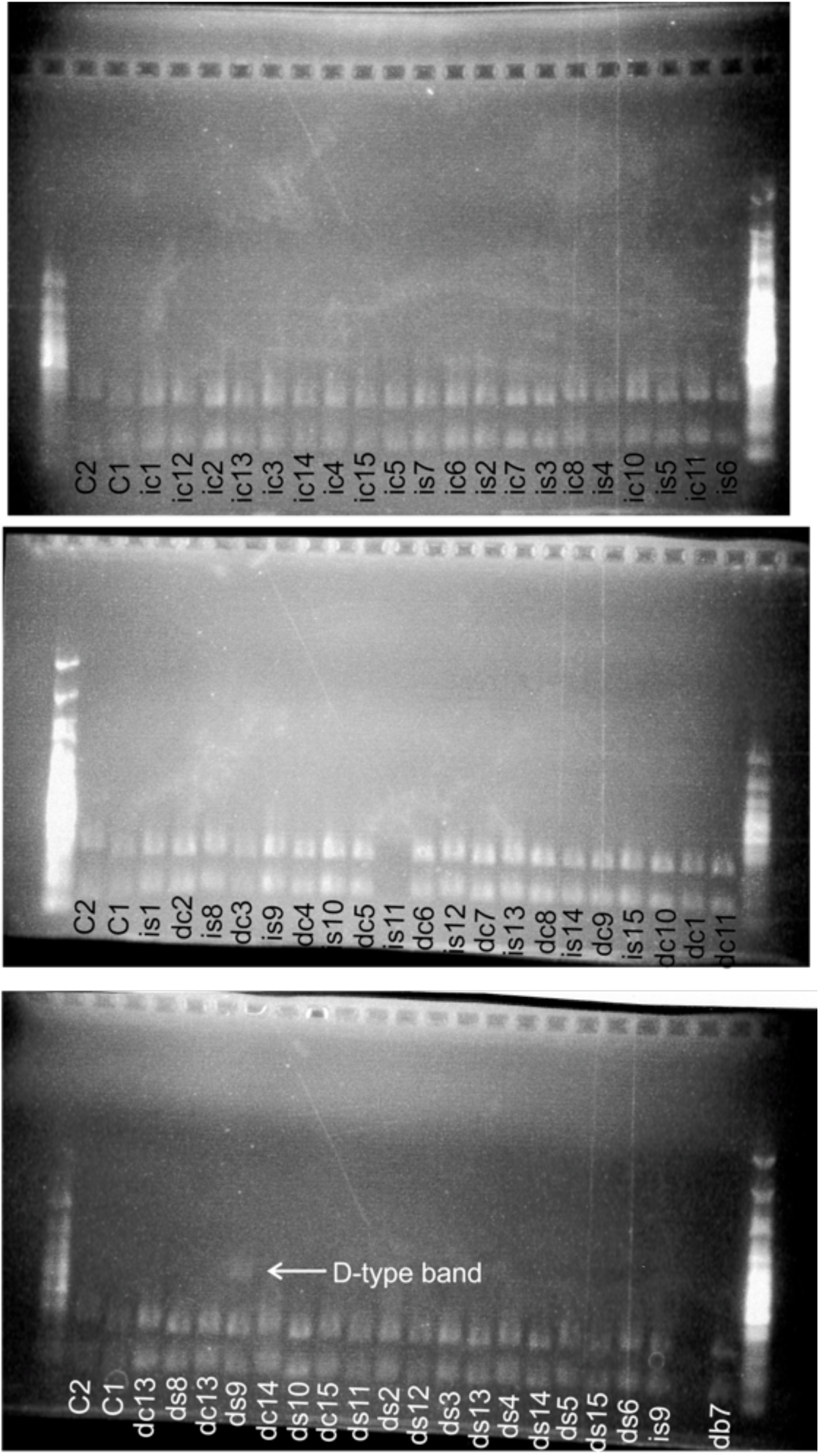
Electrophoresis gel showing digest of *Symbiodinium* lsu type for sampled corals. All banding patterns match C1, save for sample ds9. C2 = *Symbiodinium* type C2 banding pattern, C1= *Symbiodinium* type C1 banding pattern. ic=Upa-Upasina Control, is=Upa- Upasina Seep, dc=Dobu Control, ds=Dobu Seep.

**Figure S2.**
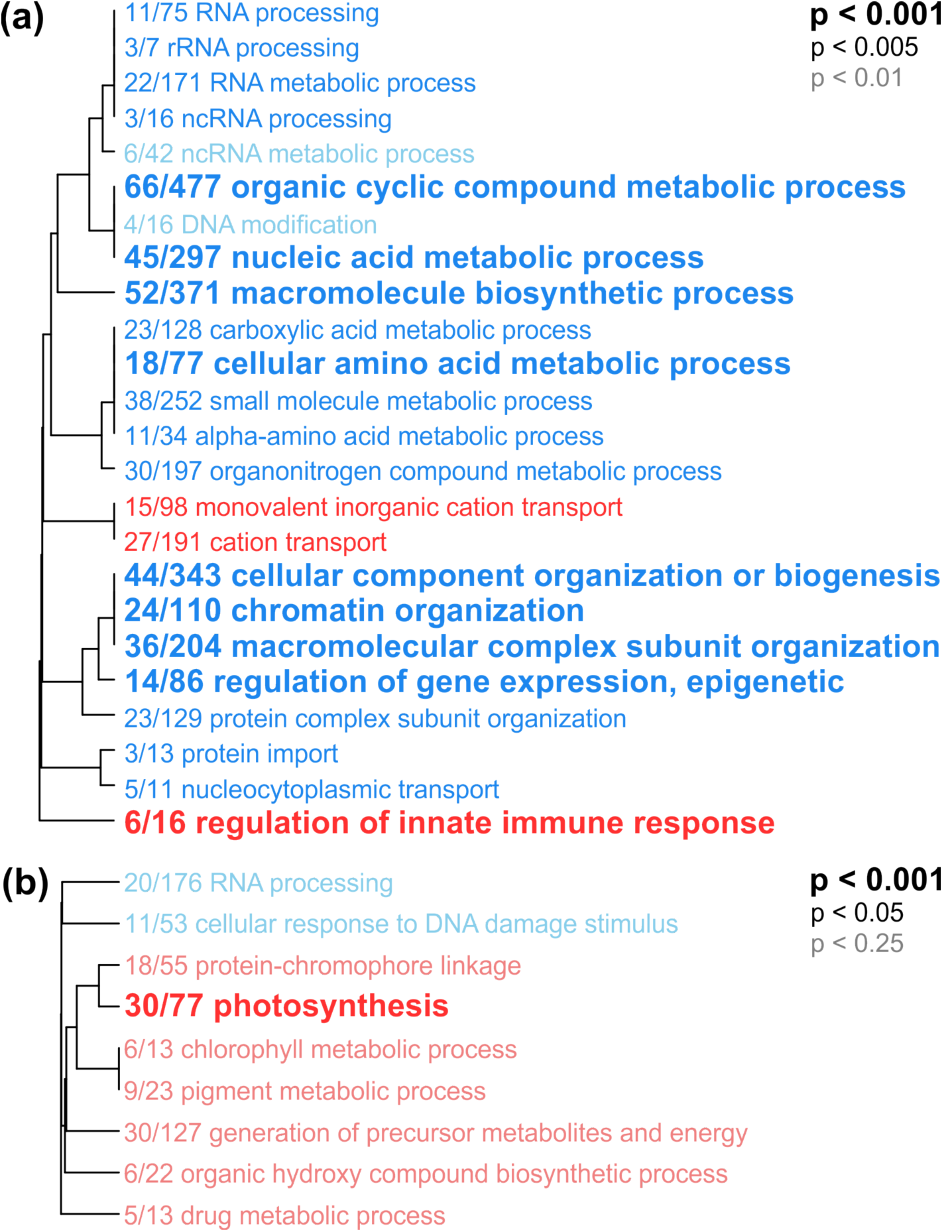
Hierarchical clustering of enriched gene ontology terms (‘biological process’) among differentially regulated genes by reef origin for the coral host (a) and symbiont (b). Red indicates terms among genes upregulated in Upa-Upasina-origin corals relative to Dobu corals and blue indicates terms among genes upregulated in Dobu corals relative to Upa- Upasina corals.

